# Cloud-Connected Patch-Worn Auscultation Device for Chest Sound Monitoring

**DOI:** 10.1101/2024.07.03.601965

**Authors:** R. Scott Downen, Baichen Li, Quan Dong, Zhenyu Li

## Abstract

In recent years, wearable electrocardiograms have risen in popularity as a solution for personal monitoring of heart activity. However, this technology has limitations in diagnostic capability and structural function monitoring. Meanwhile, auscultation of the heart remains a fundamental tool for physicians in diagnosis and monitoring of heart disease largely unaddressed in a convenient wearable format. The present work outlines a promising system currently under investigation, allowing user-initiated 10-second chest-sound recordings to be transmitted over Bluetooth-Low-Energy, with an innovative package design providing inherent noise reduction and a high signal-to-noise ratio. The device has been tested on healthy individuals, and system response has been validated against calibrated electrocardiogram recording equipment to analyze signal capture fidelity.

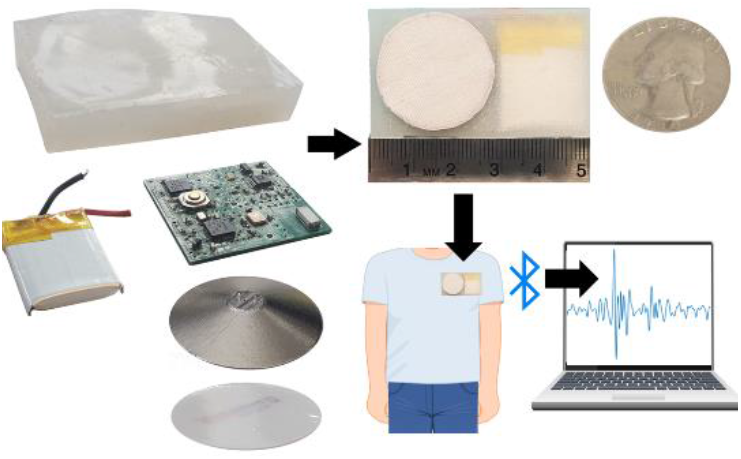

## I. Introduction

BETWEEN 1900 and 1960, life expectancy increased from 47.3 years to 69.7 years, largely attributed to sanitation norms, antibiotics, and the development of vaccines. However, this improvement in sanitation led to heart disease, cancer, and strokes replacing infectious diseases as the leading cause of death [4]. During this transition phase, heart disease became the leading cause of death in the United States, and has maintained this position for 100 years [1]–[3]. Today, of the 805,000 Americans who have a heart attack every year, 4 of 5 is symptomatic, and 200,000 have previously had a heart attack [5]. Additionally, in 2009-2010, 46.5% of US adults over the age of 19 had at least one risk factor for cardiovascular disease (CVD) and stroke [5]. Although age-adjusted death rates from CVD have declined since 1968, partially thanks to progress in prevention and treatment [3], patient outcomes following major heart events remain bleak [6]. A survey of 500 heart attack survivors indicated that 1 in 4 patients do not return to work, less than 33% understand the nature of their condition, and a large number face difficulties with every day activities. This difficulty has been attributed to a lack of awareness and understanding of their condition and recovery, leading to survivors who struggle to return to their pre-heart event lifestyles, thus increasing the risk of future heart events [6].

Providing a convenient cardiac monitoring system for heart attack survivors may be the missing piece to the puzzle in helping them return to pre-heart event lifestyles. While considerable progress has been made in the field of wearable heart monitoring thanks to the advent of miniaturized electrocardiographic (ECG) methods, the ECG, which detects electrical signals, may miss structural irregularities which are more likely to present as mechanical vibrations detectable through auscultation [2], [7]. More specifically, abnormalities in natural or implanted valves or murmurs which may be indicative of pathological cardiac defects including heart failure, arrhythmia and cardiomyopathy are difficult to observe through an ECG [1], [2]. Heart auscultation remains a fundamental tool in the diagnosis and monitoring of heart disease largely unaddressed in a wearable format [1].

Concurrently, despite being one of the traditional diagnostic tools utilized in monitoring heart conditions, stethoscopes have been declining in popularity [8]. This reduction in use may be explained by a decline in auscultatory proficiency of physicians due to a lack of skilled teachers, reliance on more elaborate and expensive tools, and a volume-driven reimbursement system that does not reward time spent with patients [8]. Additionally, physician cardiac examination skills have been observed to decline after years in practice [9]. It is believed the electronic stethoscope may help alleviate this decline, with amplification and filtration methods increasing murmur detection sensitivity [10]. Further, digitally acquired heart sounds may be combined with machine learning techniques and Artificial Intelligence to assist physicians with diagnoses of conditions such as pulmonary hypertension [11]. In this sense, the emerging field “computer aided auscultation” may represent a tremendous opportunity for the medical community [12].

Recent research demonstrated that the diagnostic capabilities of a stethoscope, when used in less traditional means, remains high. For example, the phonocardiogram was created as a visual representation of sounds collected through the stethoscope as a waveform display [13]. Mathematical methods have been shown to improve diagnostic capabilities using phonocardiograms by analyzing these waveforms using time, frequency, and energy information [14]. In fact, in some cases results from phonocardiogram analysis has been used as a gold standard to assess the diagnostic capabilities of physicians [15]. Still, intelligent auscultation technology is not widely used to assist in clinical diagnosis [16], and better tools for research and application of computer-aided heart sound analysis are needed to grow this field [12]. In addition to this need, simple and automated screening methods for valvular heart disease would prove invaluable in low- and middle-income countries, where screening campaigns are usually limited to recognizing patients with advanced phases of heart failure [17].

Traditional analog stethoscopes amplify chest sounds to reach 60 to 75dB, making them audible to the user [18], however heart murmurs and other irregularities become difficult to hear in emergency departments where ambient noise levels regularly reach levels between 60 and 70dB [19]. Auscultation can also be impacted through the physician’s handling of the stethoscope via noise introduced by finger movement along the stethoscope or hand tremors [20]; converting the electronic stethoscope to a wearable format eliminates the impact of a physician’s contact.

The present work demonstrates one such system, a wearable personalized auscultation device. The device was designed and fabricated using a custom bell-shape design for mechanical signal amplification; a custom printed circuit board for signal filtration, amplification, collection, data analysis, storage, and transfer via Bluetooth Low-Energy (BLE); a rechargeable lithium-ion-polymer (LiPo) battery to power the system and a custom polydimethylsiloxane (PDMS) enclosure.

## II. System Design

A novel audio collection platform was designed with several key innovations outlined below. In the current form of the device, a user is able to initiate a 10-second audio sample by pressing a micro-tactile switch and is notified of active recordings through a solid blue Light-Emitting-Diode (LED). Upon recording completion, the blue LED turns off and the data is stored to on-board flash memory capable of saving up to one hundred 10-second audio samples. Lastly, all data stored on the device can be transmitted via BLE upon initiation by the central station, which may be a raspberry pi or similar Linux-based platform. Data transfer via BLE is indicated to the end user through a blinking blue LED. When not in use, the device can be placed on a charging station to recharge the attached 250mAh lithium-ion battery through gold contact pads. The battery charge status is indicated through red and green LEDs. When not actively recording or transferring data, the device firmware sets the device to a low power state to maximize battery life. Future firmware will be designed to further stretch battery life; however, these efforts will rely heavily on algorithm development and the capability of the device to detect chest-sound abnormalities to initiate recordings, a process which relies on controlled clinical recordings being pursued with the current device. A more detailed description of the fabricated device subsystems shown in Fig. 1 is provided in the following subsections.

**Fig. 1.**
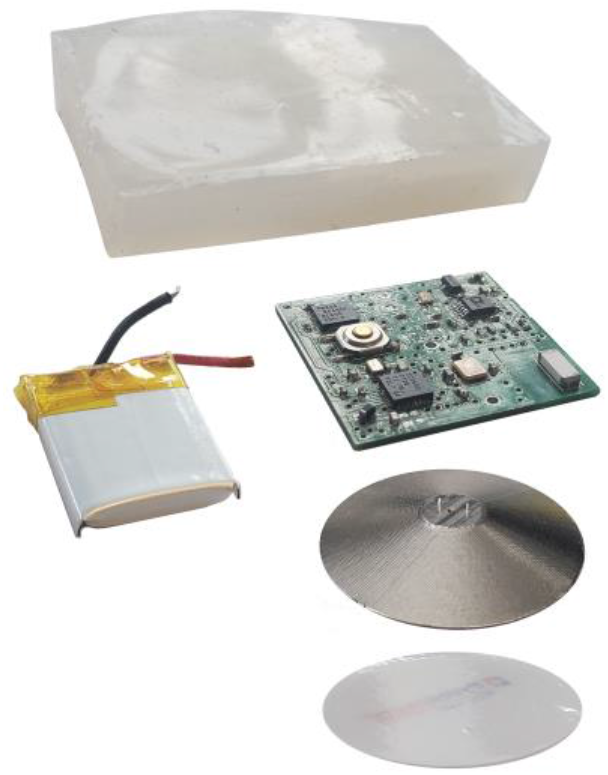
Exploded view of patch components. A custom PDMS case houses the rechargeable LiPo battery, custom circuit board assembly, and custom aluminum bell assembly.

### A. Piezoelectric Audio Collection

Several previous designs exist for wearable chest-sound audio recording, but these systems typically rely on traditional capacitive or accelerometer-based MEMS audio sampling techniques [21]–[25], which often lack mechanical noise reduction due to omni-directional audio collection. In comparison, at the center of the present circuit design is a microelectromechanical-system (MEMS) piezoelectric-based microphone (Vesper VM1000). The crux of this design, this microphone was selected based on noise isolation properties inherent to piezoelectric material, hereafter referred to as ‘piezo’, along with environmental durability provided through the MEMS device. While patient-side and outside world noise can still be transmitted to the piezo, sensitivity of piezo materials to off-axis stimuli is typically low [20], thus reducing a significant source of noise in a wearable format due to clothing or similar sheer forces applied to the patch.

The piezo sensing element helps with low-power requirements of the system compared to traditional capacitance-based microphones as it does not require an active voltage bias [26]. Further, inherent to the MEMS piezoelectric design are improved noise immunity and signal-to-noise ratios (SNR) due to higher voltage outputs converted through the nature of a piezoelectric element. Future studies may involve piezo element and chamber optimizations to improve SNR in the desired frequency range.

### B. Circuit Design and Fabrication

A block diagram of the overall system architecture is provided in Fig. 2. To support the microphone and system requirements, a custom circuit was designed for low power audio collection, storage, and secure wireless transmission, and a 1” x 1” x 0.032” printed circuit board (PCB) was designed and fabricated based on this design as shown in Fig. 3.

**Fig. 2.**
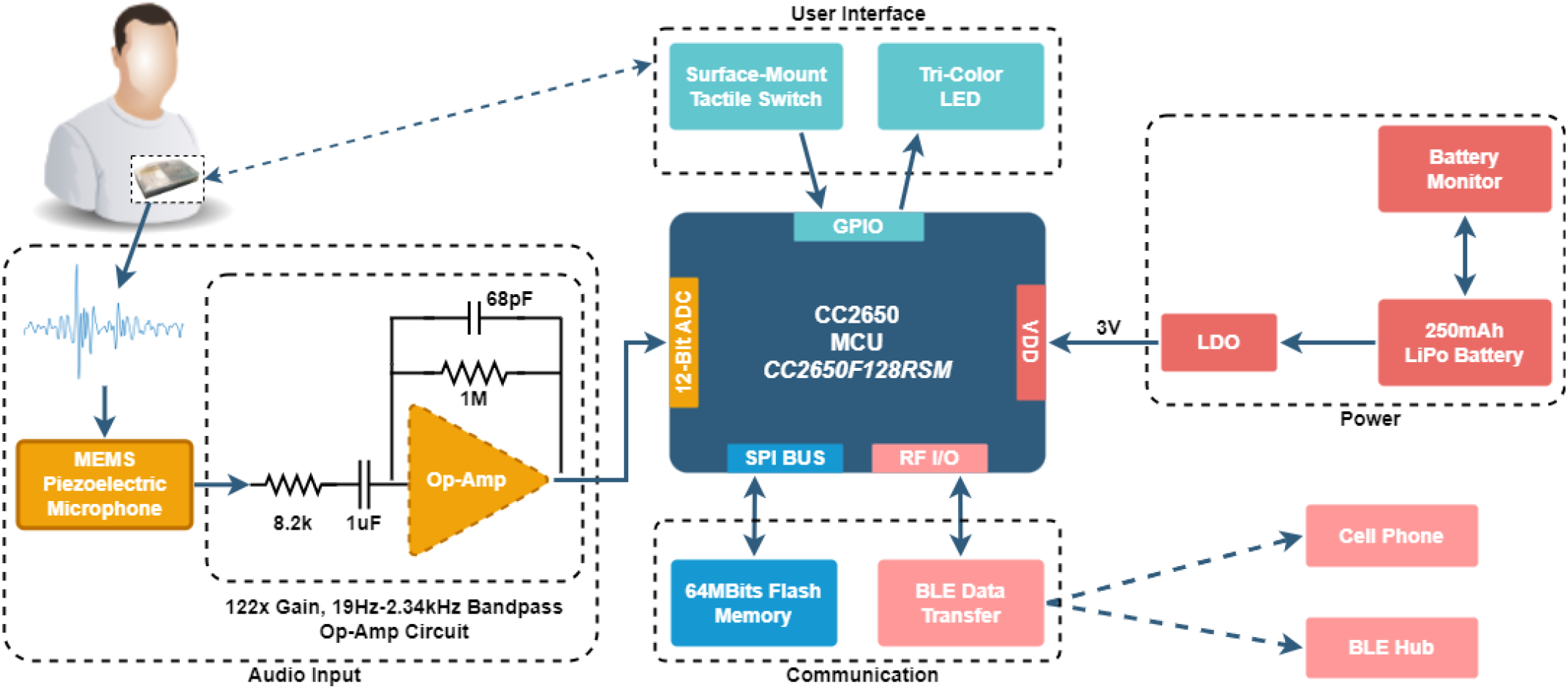
High level system architecture. Within the patch format, a tactile switch initiates audio collection via MEMS piezo microphone, which is amplified through an active bandpass circuit with 122x Gain. A 12-bit ADC on-board the CC2650 microcontroller collects the audio data, which is stored in discrete 64MBit flash memory. The system is powered by a 250mAh rechargeable LiPo battery, and user feedback is provided via tri-color LED. Upon external initiation, on-board BLE transfers stored audio to an external collection platform for further analysis.

**Fig. 3.**
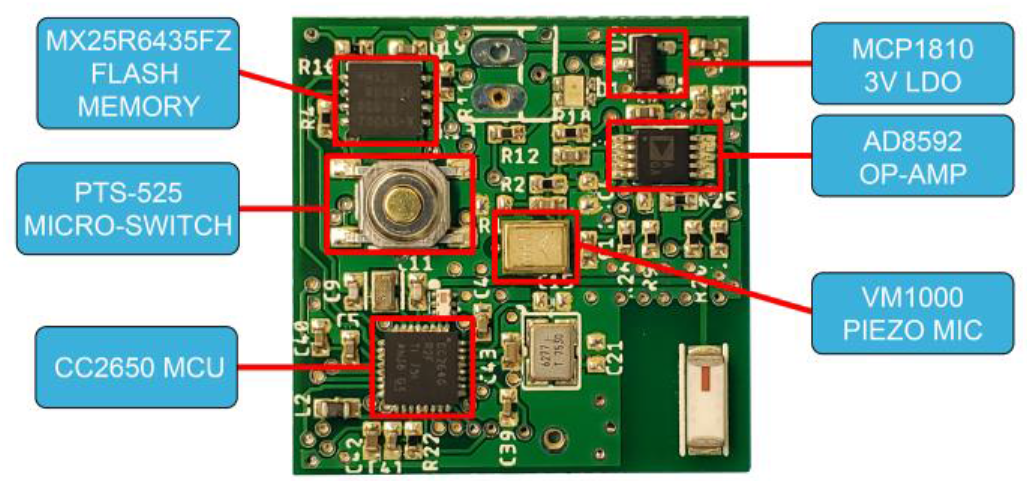
Custom auscultation device PCB assembly featuring flash memory, a low dropout linear regulator, operational amplifier to amplify and filter the incoming audio signal, a MEMS piezoelectric microphone, CC2650 microcontroller, and microswitch to initiate audio recordings. Bluetooth-low-energy is transmitted directly from the microcontroller to the ceramic antenna.

The circuit board includes a high-speed audio operational-amplifier (Analog Devices AD8592) to amplify the analog audio signal and reduce the risk of signal dropout, a 5mm X 5mm microcontroller with on-board BLE for data processing and transmission (TI CC2650), a tri-color LED and surface-mount tactile switch for user interaction, 64 M-Bit flash memory for data storage (Macronix MX25R6435FZ), a low-power 3V linear voltage regulator (Microchip MCP1810), a charge management and battery protection chip (Microchip MCP73833), and a 250 milliampere hour (mAh) rechargeable LiPo battery.

A band-pass filter was built into the amplification circuit to allow frequencies from 19Hz to 2.34kHz to target primary frequencies of the heart, which typically lie between 25Hz and 200Hz [16]. Should the device be used for other physiological indications, such as lungs sounds, this circuit can be adjusted along with the sampling frequency set in firmware or software.

### C. Bell Design

A custom aluminum stethoscope bell was designed for use with the system and fabricated using a desktop computer-numerical-control (CNC) mill (Carbide Nomad 883 Pro). For the first design, aluminum was selected for its strength-to-weight ratio and ability to be fabricated on our available CNC. In future iterations, denser metals such as titanium or stainless steel may be considered. Readily available, off-the-shelf stethoscope membranes were cut to size to use with the custom bell and glued on using Loctite Ultra Gel Control. The bell subassembly was then aligned with holes in the circuit board using designed-in alignment pins and glued to the PCB for permanent attachment. Future work is expected in simulating and optimizing bell parameters for use with the developed platform.

### D. Patch Design

When considering the ideal format for the current device, a number of form factors were considered, including the Holter-type, harnesses, bands, and sensor patch [27]. Based on the desired use of on-demand or continuous monitoring, a patch format was decided upon due to its small form factor and comfort over long-term use without external wires or straps. For heart sounds, placement of the device underneath the left pectoralis major is one of the ideal collection sites, as well as one of the preferred regions to minimize interference with human movement while preventing removal of the patch due to twisting or stretching of the skin [28]. This desired placement led the mechanical design and overall dimensions of the patch for comfort and overall fit.

To create a patch to enclose the electronic components in a wearable format, a simple mold was designed and fabricated. First, a mold was designed with cavity dimensions 0.5mm larger than those of the electronics, battery, and bell, creating the overall thickness of the enclosure walls. Next, a 3D proxy for the electronics and battery was designed as a single piece in Solidworks CAD. These representative components were 3D printed on a Zortrax M200 3D printer and attached to the bell location using a single M2 × 3mm screw as shown in Fig. 4(a). This mounting position for the two mold pieces was selected to create a hole for the bell to be mounted through during assembly. Dragonskin 10 Very Fast flexible silicone was poured into the cavity as shown in Fig. 4(c), and a wooden stir stick was used as a squeegee to create a flat top surface, Fig. 4(d). The silicone was then allowed to cure for 45 minutes at room temperature. After curing, a razor was used along the outside edge of the patch to ensure a clean removal from the mold. A small slit was then cut above the mounting screw and the screw was taken out. After removing the patch from the mold, a slit was cut along the long edge of the patch and the 3D printed insert was removed. The resulting patch is shown in Fig. 4(f).

**Fig. 4.**
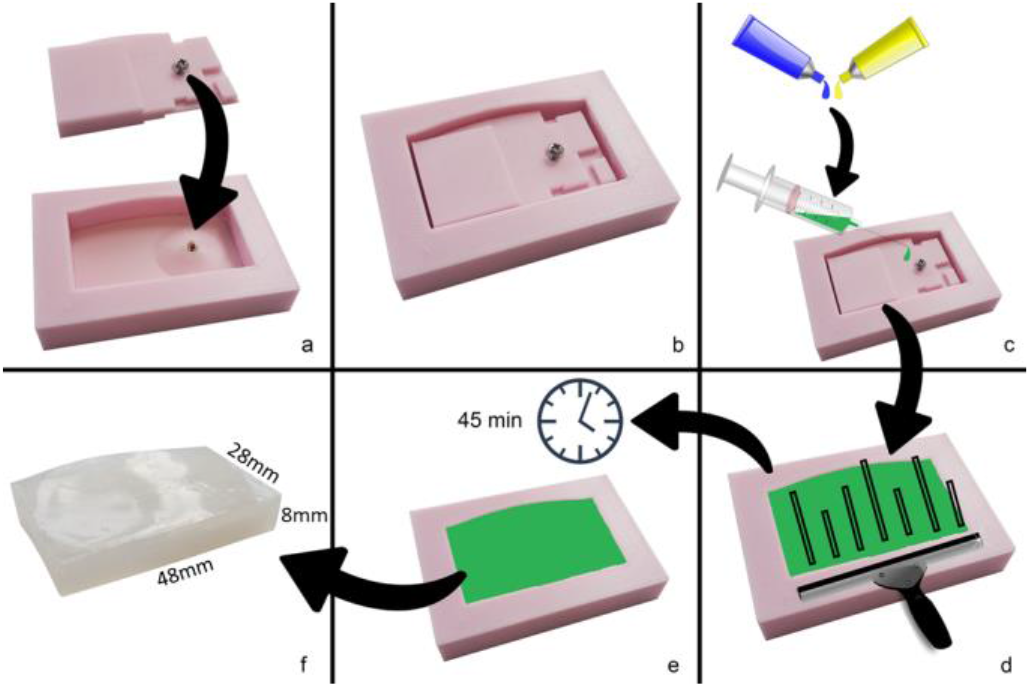
Patch fabrication. (a) 3D-printed insert PCB outline attached to outer wall cavity; (b) Final mold; (c) Silicone mixed and injected into the mold; (d) Squeegee of silicone to provide a flat curing surface; (e) 45 minute curing time; (f) Final patch removed from mold

### E. Embedded Code

To run the device, the CC2650 microcontroller was programmed through Code Composer Studio. Two primary power states were used: standby and active, with active power state broken down as BLE-active transmission, and audio collection. In the current implementation, the device stays in standby mode until a user initiates a 10-second sample using the micro tactile switch, which wakes up the device. After waking, the device begins data collection at 8kHz sampling frequency through a buffer, which pushes collected data to the on-board flash memory. Following sample collection and transfer to the flash memory, the device resumes standby awaiting another sample. In standby, the device also polls the Bluetooth channel awaiting a connection—an external device can initiate a data pull, which wakes the device and begins transferring data from the external flash, through the CC2650 (using a similar buffer scheme), and to the external Bluetooth device. Once the data transfer is complete, the external flash memory is erased automatically, and the system resumes standby.

### F. Final Assembly

After fabricating all subsystems and programming the CC2650, the final patch was assembled. First, the LiPo battery was directly soldered to the circuit board using through-hole locations in the PCB. The electronics were then slid into the patch through the slit created along the long edge. A thin layer of Loctite Ultra Gel Control was spread along the mounting surface of the fabricated bell, and it was pressed onto the PCB through the hole in the patch using through-hole alignment holes in the PCB and small bosses designed onto the bell. Following this assembly, the final overall dimensions of the patch are 48mm x 28mm x 8mm. For the initial prototype, simple medical-grade tape was used to hold the patch onto the body; in future iterations, it is expected that standard medical adhesive tapes would be pressed onto a molded-in ridge along the perimeter of the patch, similar to a picture frame.

## III. Results

Initial results for critical system parameters are provided in the sections below.

### A. Battery Life

In standby mode the device draws approximately 4µA; audio collection raises the current consumption to 5mA; lastly, when transmitting data over BLE, current consumption is approximately 11mA. As described above, for initial device collection of heart sounds, the prototype uses a user-initiated data collection method through an on-board physical switch. Based on this, assuming six samples an hour over 24-hour periods, the device is expected to last approximately 45 days before needing to be recharged. In the near-term data collection phase, one anticipated use-case transition is interval-based heart sound collection system, where the user will wear the device throughout their daily life while it collects heart-sound samples every 10 minutes. In this use case, two devices would be effective; one for daytime use, and one for night use, also encouraging the end user to change the adhesive on a regular interval to prevent skin irritation. In this case, data transmission would be automatically initiated when the device is plugged in to a charging station, thus transmission current is not factored in to calculations. In this use case, battery life is expected to last well beyond expected typical daily use below 18 hours.

Long-term, one expected use case is an interval-based abnormality monitoring system which collects an audio sample every 10 minutes, analyzes the sample, and transmits the data if any abnormalities are detected. Assuming 10-second samples are collected, and a full ten second analysis of data is performed on the device with a similar current consumption of BLE transmission, battery life is calculated to be 23 days. While the battery size could be reduced, the current form factor thus far is adequate for data collection without impeding skin contact, and future real-time data analysis and on-board algorithms based on this long-term goal may require significantly more power.

### B. Preliminary System Validation

To validate the essential performance of the wearable audio collection device, three independent tests were performed to examine signal quality and data transfer. Similar testing as that which would be performed on a digital stethoscope is appropriate to ensure proper audio capture and processing techniques, as the current device meets the Federal Drug Administration (FDA) definition of a digital stethoscope. First, the electrical system’s ability to capture, store, recall, and transfer analog data over BLE was evaluated. Second, the device’s electrical noise was quantified. Lastly, the device was compared to an FDA-cleared electrocardiogram system to validate its ability to adequately collect heart sounds. Overall system characteristics are shown in Table 1.

**TABLE 1.** System Specifications.

| Parameter | Value |
| --- | --- |
| Power Consumption – Sleeping | 4 μA |
| Power Consumption – Recording | 5 mA |
| Power Consumption – BLE Transmission | 11 mA |
| Expected Battery Life (with 240 mAh LiPo) | 45 days |
| Sampling Frequency | 8kHz |
| Bandpass Filter | 12Hz – 2kHz |
| Audio Gain | 120x |
| Storage Capacity | 100 x 10s |

<sup>a</sup>A = ampere.

#### 1) Analog-to-Digital Converter (ADC) Signal Quality

To examine the initial signal received by the circuit board, input into the microcontroller, saved to on-board flash memory, and transmitted over BLE, a simple test of the ADC was performed. This test also functioned to validate the system sampling frequency as well as the firmware loaded into the microcontroller. A signal generator set to output a 1kHz sin wave was connected directly to the input of the ADC (bypassing the microphone) to be passed through to the microcontroller, and the input switch was activated to take a 10-second sample. The audio sample was then transmitted over BLE to a computer and loaded into Matlab as a comma separated value (.csv) file. Sampling errors in the system, whether through the ADC capture, data storage, data retrieval, or transmission through Bluetooth would be detectable in the form of aliasing or signal offset observed in the power spectrum of the transmitted waveform. Observed results are shown in Fig. 5.

**Fig. 5.**
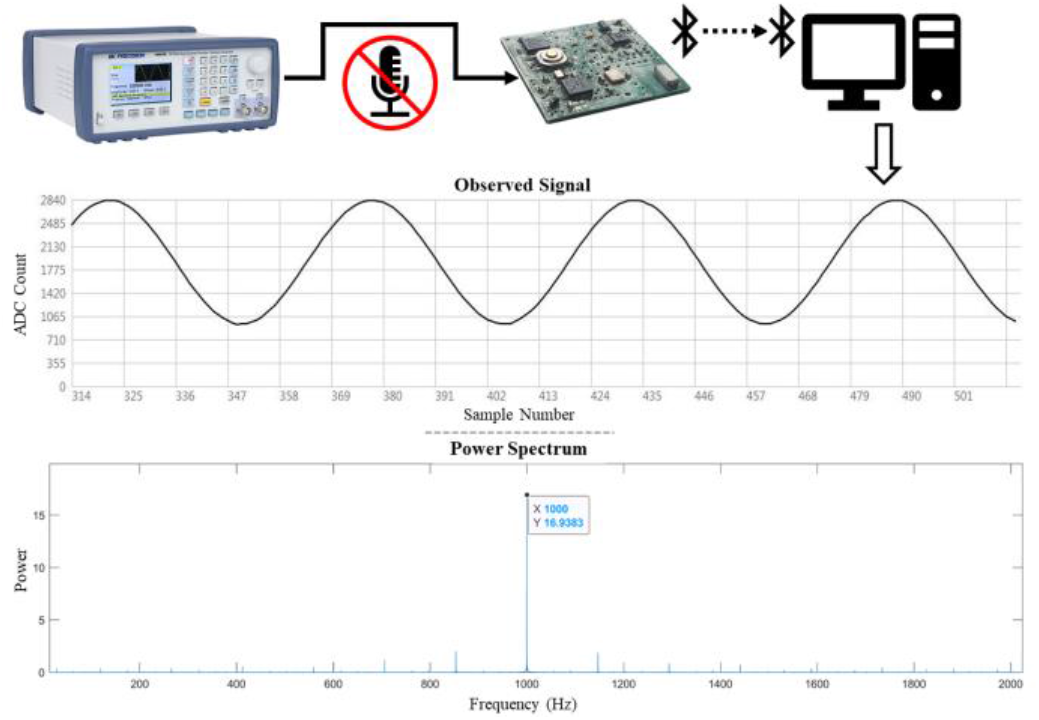
1kHz input signal observed after Bluetooth transmission to validate collection and transmission quality. Signal shows strong power at 1kHz with minimal aliasing and no frequency shift, indicating adequate data storage and retrieval, and no transmission loss.

#### 2) Device Electrical Noise

As is common in sensor devices, the electronic noise floor for the system was analyzed. The simplest method of analyzing noise generated by the electronic components and PCB design was accomplished by removing the piezo device from the system (by removing a series resistor in the circuit, thus disconnecting the MEMS piezo device). After recording a 10-second sample and transmitting over BLE, the ADC signal can be converted to volts. Following this process, the average electronic noise floor was determined to be approximately 6 mV, shown in Equations (1) and (2).

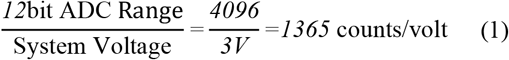

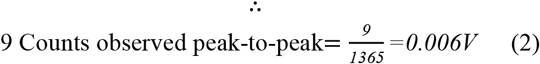

#### 3) Device Comparison with ECG Machine

Following the computational analysis provided by Ayazi et. al [29], the heart sounds collected through the wearable device were compared with ECG signals collected through a calibrated benchtop ECG machine, as shown in Fig. 6(a). By extracting inter-beat intervals (IBIs), a direct comparison could be made between the collected sounds and electrical activity of the heart. Opening of the aortic valve was used as an alignment point of reference to the R-peak of the ECG. After aligning the signals, the IBIs were calculated using linear fit with r^2^ as shown in Fig. 6(b), showing a correlation of 0.9997. Lastly, a Bland-Altman plot was generated to compare the calibrated ECG data against the auscultation patch under test, which demonstrated a 95% confidence interval having a range <0.01s, as shown in Fig. 6(c).

**Fig. 6.**
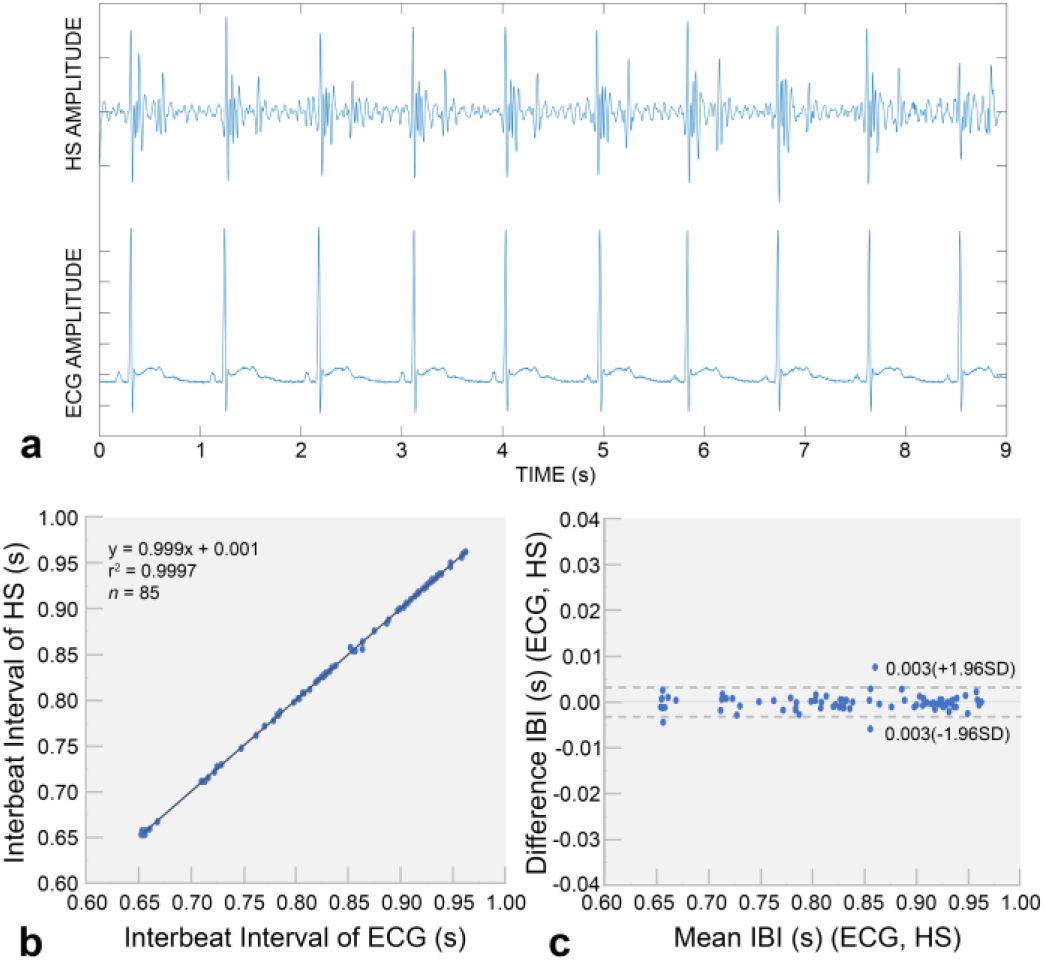
Calibrated ECG vs. auscultation patch heart interbeat interval comparison. (a) Time domain plot of ECG alongside auscultation patch data; (b) Linear curve fit of correlation plot showing r^2^ = 0.9997; (c) Bland-Altman plot comparing the calibrated ECG to the auscultation patch under test demonstrating 95% confidence interval having a range <0.01s.

## IV. Conclusion

This article presents an innovative design for a wearable auscultation device currently under investigation for heart and lung sound abnormality detection. Following initial device design, the overall system was tested for audio fidelity, ADC conversion, wireless data transfer, and amplifier performance first by injecting a waveform into the audio circuit directly, then by removing the piezo device from the circuit and measuring the electronic noise floor. Lastly, by taping the device to the chest of an adult male, the device was evaluated against a calibrated benchtop ECG machine for beat-to-beat interval alignment, showing a strong correlation, indicating adequate signal collection.

In the immediate future, we anticipate this device being used for data collection in two clinics: first, in an emergency department to collect data from healthy and affected hearts; second, in a severe pediatric asthma clinic to collect lung sounds in a controlled environment after adjusting the signal bandpass filter to accommodate lung sounds. This evaluation period will allow a greater understanding of heart and lung sounds as collected through the device as well as any limitations the current form factor, collection method, and ambient isolation methods may create. Following this data collection and performance evaluation period, algorithms for on-board automated abnormality detection will be evaluated and compared against audio collected through a commercially available digital stethoscope as evaluated by trained physicians. Long-term, we believe this work will provide a data collection platform to enable future studies involving trained models in real-world environments with the ultimate goal of providing a monitoring and early alert system for individuals with heart and lung conditions.

---

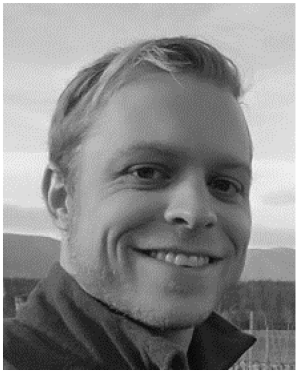

**R. Scott Downen** R. Scott Downen received a B.S. degree in Applied Sciences with a focus in Biomedical Engineering from the University of North Carolina, Chapel Hill, in 2011 and a M.S. degree in Biomedical Engineering from The George Washington University, Washington, DC, (GWU) in 2017. In 2023, he received a Ph.D. in Biomedical Engineering from GWU with a focus in microfluidics and point of care molecular diagnostics platforms. His research interests include, but are not limited to, wearable sensors, microfluidics, rehabilitative technology, and embedded and computer-aided design.

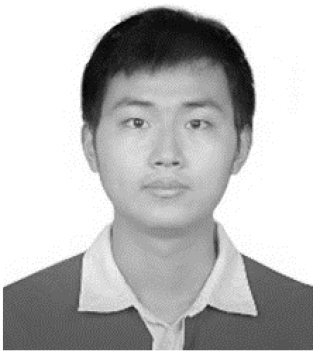

**Baichen Li** received a B.S. degree from The Hong Kong Polytechnic University in 2009, a M.S in Electrical and Electronics Engineering from The University of Hong Kong in 2010, and an M.S. in Biomedical Engineering from the George Washington University (GWU) 2013. Following this, he worked in industry as an electrical engineer before returning to GWU, where he received a Ph.D. in Biomedical Engineering in 2021 with a focus on wearable sensors, wireless sensor networks, point-of-care diagnostics, and microfluidics. Following his Ph.D., Baichen returned to China to work as a post-doc in China Investment Corporation (CIC) with a focus in healthcare and biotechnology..

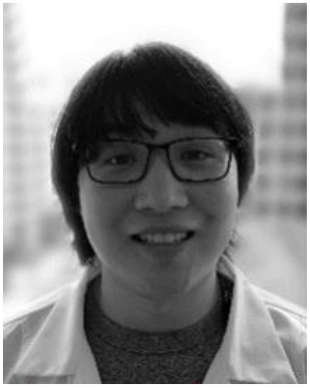

Quan Dong received a B.S. degree in Automation from the Beijing Institute of Technology, Beijing, China in 2011, and a PhD degree in Biomedical Engineering from the George Washington University, Washington D.C, USA, in 2021 with a focus in wearable sensor networks and microfluidics. He is now a staff systems engineer in Samsung Research America working on wearable medical devices. His areas of interest include biomedical devices, wearable devices, and flexible electronics.

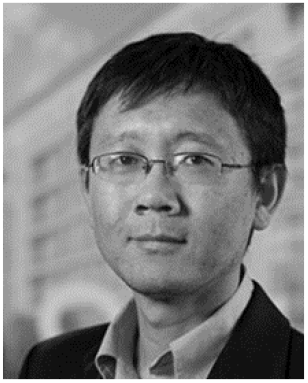

Zhenyu Li (M’08) received the B.S. degree in precision instruments from Tsinghua University, China, in 1999, the M.S degree in electrical engineering from University of California at Santa Barbara, CA, in 2000, and the Ph.D. degree in electrical engineering from the California Institute of Technology (Caltech), Pasadena, CA, in 2008. From 2008 to 2010, he was a postdoctoral scholar at the Howard Hughes Medical Institute Janelia Research Campus and Caltech. Now he is Associate Professor with the Biomedical Engineering Department, The George Washington University, Washington DC. His current research interests include the development of biosensors and medical devices using microfluidics, MEMS, Optofluidics and flexible electronics. He has published over 50 peer-reviewed articles and holds seven patents.

